# Cytokine-expression patterns reveal coordinated immunological programs associated with persistent MRSA bacteremia

**DOI:** 10.1101/2022.12.28.521386

**Authors:** Jackson L. Chin, Zhixin Cyrillus Tan, Liana C. Chan, Felicia Ruffin, Rajesh Parmar, Richard Ahn, Scott Taylor, Arnold S. Bayer, Alexander Hoffmann, Vance G. Fowler, Elaine F. Reed, Michael R. Yeaman, Aaron S. Meyer, the MRSA Systems Immunobiology Group

## Abstract

Methicillin-resistant *Staphylococcus aureus* (MRSA) bacteremia is a common, life-threatening infection that imposes up to 30% mortality even when appropriate therapy is used. Despite *in vitro* efficacy, antibiotics often fail to resolve the infection *in vivo*, resulting in persistent MRSA bacteremia. Recently, several genetic, epigenetic, and proteomic correlates of persistent outcomes have been identified. However, the extent to which single variables or composite patterns operate as independent predictors of outcome or reflect shared underlying mechanisms of persistence is unknown. To explore this question, we employed a tensor-based integration of host transcriptional and proteomic data across a well-characterized cohort of patients with persistent and resolving MRSA bacteremia outcomes. Tensor-based data integration yielded high correlative accuracy with persistence and revealed immunologic signatures shared across both the transcriptomic and proteomic datasets. We find that elevated proliferation of mature granulocytes associates with resolving bacteremia outcomes. In contrast, patients with persistent bacteremia heterogeneously exhibit correlates of granulocyte dysfunction or immature granulocyte proliferation. Collectively, these results suggest that transcriptional and proteomic correlates of persistent versus resolving bacteremia outcomes are complex and may not be disclosed by conventional modeling. However, a tensor-based integration approach can help to reveal consensus molecular mechanisms in an interpretable manner.

**Significance Statement:** While antibacterial therapies effectively resolve MRSA *in vitro*, these treatments often fail to clear MRSA bacteremia *in vivo*, suggesting that host-pathogen interactions are essential to persistent MRSA bacteremia. Recent studies have identified genetic, transcriptomic, and proteomic determinants of MRSA persistence. These determinants independently, however, provide insufficient mechanistic insight and it is unclear if they indicate unique or overlapping persistence mechanisms. Here, we use tensor-based decomposition to jointly analyze cytokine and transcriptomic measurements from patients with MRSA bacteremia. Results indicate that persistence mechanisms integrated across biological modalities reflect diverging mechanisms of persistent bacteremia. Ultimately, these results may help to identify future therapeutic targets for treating persistent MRSA bacteremia.

## Introduction

Methicillin-resistant *Staphylococcus aureus* (MRSA) bacteremia is a common, life-threatening infection, arising through both community-acquired and healthcare-associated settings (1, 2). These infections are associated with poor outcomes, and up to 30% of appropriate antibiotic regimens fail to resolve bacteremia *in vivo* despite efficacy *in vitro* (3). MRSA bacteremia that resolves upon appropriate antibiotic treatment is termed resolving bacteremia (RB) whereas cases that do not resolve after 5–7 days of therapy are termed persistent bacteremia (PB) (4). The limited predictive value of *in vitro* susceptibility for MRSA bloodstream clearance clinically indicates a need to better understand the determinants of antibiotic therapy outcomes *in vivo*.

Recent progress has been made in identifying determinants of PB vs. RB outcomes in MRSA bacteremia (4–6). Host factors appear to play an important role in MRSA persistence, as patient outcome can be independent of strain susceptibility to vancomycin or daptomycin susceptibility *in vitro* (3). Persistence factors are distinct from those associated with MRSA antibiotic resistance, where the organism is refractory to the antibiotic both *in vitro* and *in vivo* (7). Thus, advances are necessary to better discern and predict persistence outcomes *in vivo*. We hypothesize that outcomes are determined by the confluence of immunological responses in an individual host, the infecting MRSA strain, and the specific antibiotic and its use practice.

To this end, we have previously undertaken broad molecular profiling to measure the molecular differences of MRSA bacteremia response (4–6). These systems-level analyses have identified genetic (5), transcriptional, and cytokine (6, 8) correlates of MRSA bacteremia persistence outcomes. However, the extent to which these signatures operate as distinct molecular mechanisms of phenotypic immune response or reflect a shared underlying immune program is yet unclear. Therefore, we approached this problem based on the premise that shared patterns of molecular and cellular responses might improve understanding of clinical correlates of outcome if they each reflect integrated molecular mechanisms.

Matrix and tensor factorization techniques are powerful tools for reducing the dimensionality of complex data. Most generally, these methods reduce multi-modal datasets (data that can be arranged into several dimensions, such as measurements, patients, and time) into mode-specific matrices that individually capture patterns across each dimension. These factor matrices individually reveal unforeseen relationships among the diverse datasets and, when recombined, approximate the original measurements. These methods, when appropriately matched to the structure of the data, help to visualize its variation, reduce noise, impute missing values, and reduce dimensionality (9). For data in matrix form, principal components analysis (PCA) and non-negative matrix factorization are two examples widely applied (10). When integrating data of higher dimensions, higher-order generalizations of these methods, tensor factorizations, can be applied (9).

A particularly important benefit of factorizing data into mode-specific matrices is that it is naturally suited to combining different sources of data which often derive from diverse biological measurements. Variation along each mode of the data in tensor form is effectively separated by these techniques (11, 12). When integrating two sources of data each in a matrix or tensor format, coupled matrix-tensor factorization allows one to detect shared patterns between datasets of differing dimensionality (11–13). Recognizing coupling across datasets provides two distinct benefits: (1) the extent of data reduction is increased by using a common set of patterns across both datasets; and (2) patterns distinguished in the shared mode reflect the trends presented in both datasets, thus their definition is better shaped. Consequently, retrospective associations or prospective predictions based on these factorized patterns may be improved through more accurate derivation, and interpretation of the resulting patterns may be improved by a broader, holistic view of the measurements tied to those patterns (11, 14).

In the present study, we applied coupled matrix-tensor factorization to integrate the transcriptional and cytokine responses relative to persistent vs. resolving clinical outcomes in MRSA bacteremia. Data integration enabled the identification of consistent patterns of immunologic response across both data sources and revealed patterns distinguishing PB from RB outcomes. The combined immunologic patterns explain outcome better than either data type on its own, with correlative accuracy verified in an independent cohort. These associations are shaped by two granulocyte formation and activation patterns that retain their correlational accuracy when used alone. Overall, the current results demonstrate that robust correlative relationships detected by tensor-based modeling reveal integrative immunological signatures of persistent vs. resolving outcomes in human MRSA bacteremia.

## Results

### A tensor-based strategy for integrating heterogeneous clinical measurements

To identify common patterns across cytokine and RNA-seq measurements, we first sought to optimally organize the multiple datasets. Measurements were of three types—plasma cytokines, serum cytokines, and RNA-seq from whole blood samples (Figure 1A). Known differences exist in cytokine measurements between plasma and serum; however, certain shared variation across patients is expected and thus cytokine measurements across serum and plasma sources could be aligned. Therefore, while not every patient had every type of measurement, the study contained two types of measurements that could be aligned across cytokines and patients, and RNA-seq measurements that only share the patient dimension.

**Figure 1.**
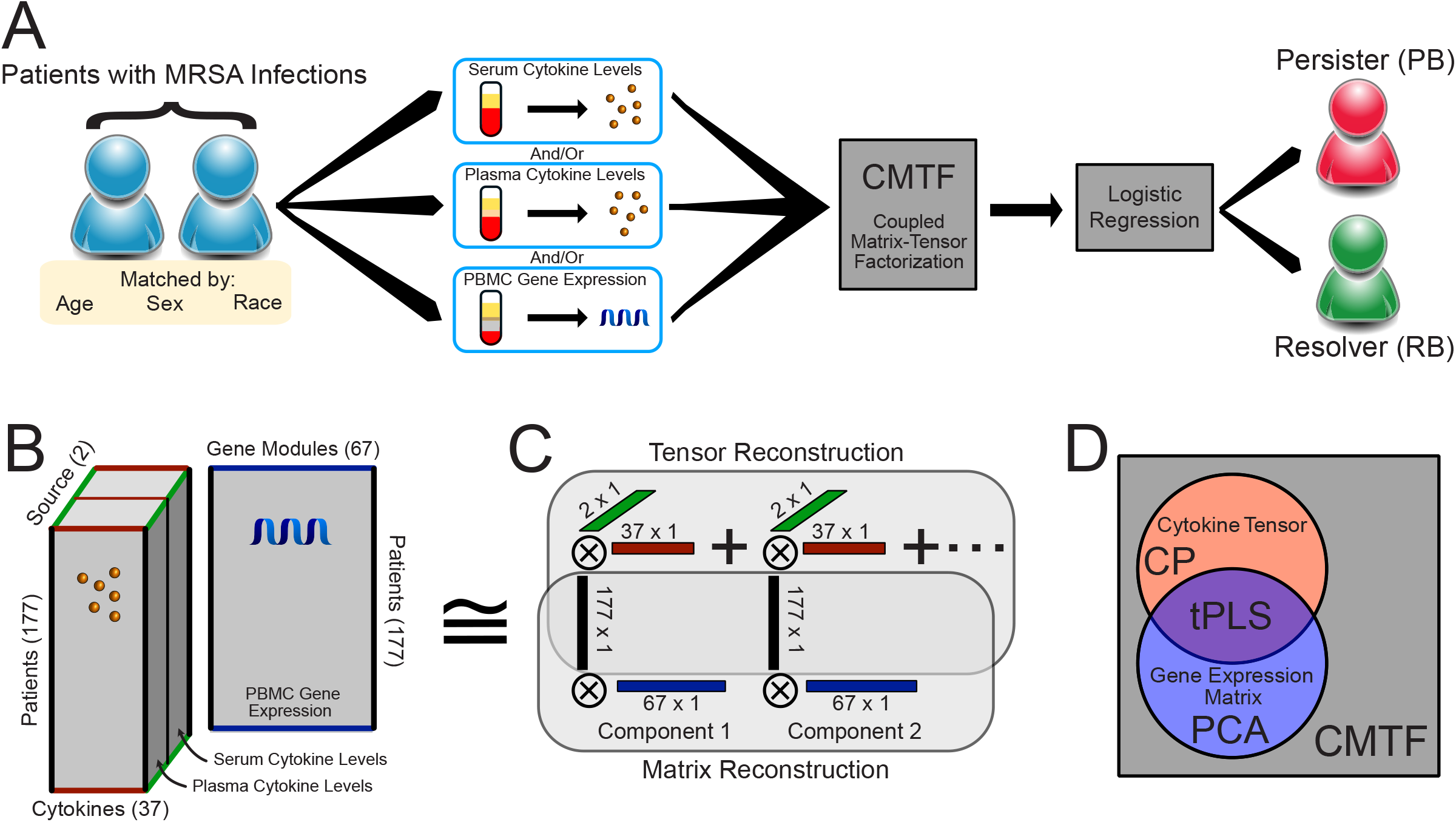
Structured data decomposition integrates clinical measurements with varying degrees of overlap. A) General approach description. Patients with MRSA bacteremia treated with vancomycin had samples collected at admission and then were monitored 5 days post-admission for clearance of MRSA from the bloodstream. Measurements of patient serum cytokine, plasma cytokine, and whole blood transcriptional profiles were assessed. These measurements were then reduced into overall factors describing patterns within the data, which in turn were used to assign disease outcomes, defined as resolving (RB) or persisting (PB) bacteremia. B) Overall structure of the data. Cytokine measurements from either plasma or serum can be arranged in a threedimensional tensor, wherein each dimension indicates patient, cytokine, or sample source, respectively. In parallel, gene expression measurements are aligned with cytokine measurements by virtue of sharing patients. C) Data reduction is performed by identifying additively separable components represented by the outer product of vectors along each dimension. The patient factors are shared across both the tensor and matrix reconstruction. D) Venn diagram of the variance explained by each factorization method. Canonical polyadic (CP) decomposition can explain the variation present within the cytokines tensor, or principal component analysis (PCA) could be used to reduce the gene expression matrix (9). Tensor partial least squares regression (tPLS) allows one to explain the shared variation between the matrix and tensor (33, 34). In contrast, here we wish to explain the total variation across both the tensor and matrix. This is accomplished with CMTF (11–13).

In situations where measurements can be aligned along two or more dimensions, measurements can be structured into tensor form. A generalization of a matrix, which is a two-mode tensor, tensors most often refer to organized arrays with three or more dimensions (modes); in the three-mode form, tensors are structured as a cube of measurements. We started by structuring the cytokine data into a 3-mode tensor, with patient, cytokine, and cytokine source (serum or plasma) axes (Figure 1B). This tensor was paired with the RNA-seq measurements in a matrix (2-mode tensor) containing the shared patient axis and a separated gene axis. We additionally summarized the RNA-seq measurements into co-expressed gene modules to reduce model complexity (15). To integrate both data types, we used coupled matrix-tensor factorization (CMTF). This method solves an optimal low-rank approximation of both datasets while keeping the coupled dimension (in this case, patients) shared during the process using an alternating least squares strategy (Figure 1C). As CMTF maximizes the variance explained across both datasets, it is better suited here than other approaches that maximize explanation of other subsets of variance, including canonical polyadic (CP) decomposition for just the cytokine tensor, principal components analysis (PCA) for just the RNA-seq matrix, or partial least squares regression in tensor form (tPLS) to specifically examine the shared variance (Figure 1D).

### Tuning dimensionality reduction for accurate correlations in MRSA bacteremia outcomes

Dimensionality reduction via CMTF introduces two method parameters that influence the resulting decomposition. First, decomposition can be performed using a varying number of components. We observed that a small number of components could effectively explain the data variation, with 8 components capturing >70% of the total variance while reducing the data to 16% of its original size (2192 factor values versus 14132 non-missing measurements; Figure 2A). Second, as CMTF aims to maximize the total variance of both datasets explained, the relative numerical scale between cytokine and RNA-seq measurements affects the goodness of fit for each individual dataset as CMTF will prioritize explaining patterns in the dataset with the larger scale. To explore the effect of this scaling, we tested a variety of scales; the resulting factors were responsive to the relative scale of each data type and increasing the emphasis of cytokine measurements improved the overall variance explained (Figure 2B).

**Figure 2.**
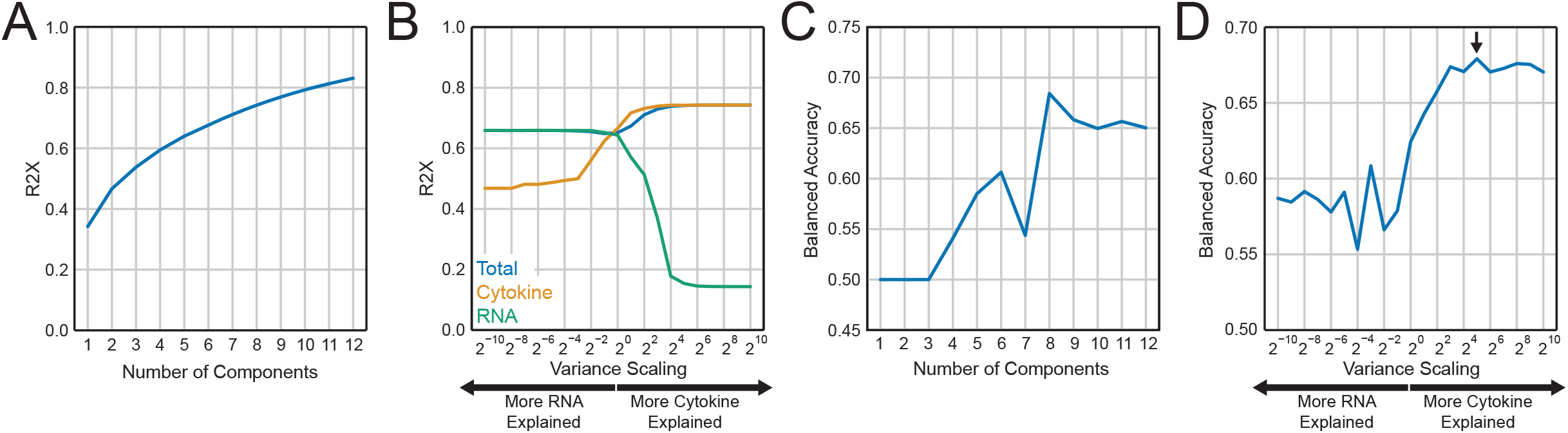
CMTF parameter tuning to correlate bacteremia outcome. A) Number of components used in the CMTF decomposition versus the percent variance reconstructed (R2X). B) Percent variance explained upon reconstruction (R2X) of the entire data, RNA, or cytokine measurements with varying scaling between the two datasets. C–D) Balanced accuracy in assigning bacteremia outcomes with varying number of components (C) and scaling (D).

We chose to identify reduced patterns that were optimally able to correctly assign PB vs. RB outcomes. To do so, we used clinical variables to correlate outcomes by logistic regression. Assignment accuracy was quantified using 10-fold cross-validation. Briefly, 10% of the patients were left out from the logistic regression model, and the remaining 90% were used to learn the relationship between each component and outcome. Next, the logistic regression model was used to assign outcomes for patients held out of the model training. This process is repeated until every patient was categorized with respect to PB vs. RB based on cytokine and transcriptome profiles. Accuracies reported are the balanced accuracy scores observed over this cross-validation process. Using this process while varying settings within CMTF, we observed a peak correlation performance at 8 components (Figure 2C) and when the cytokine data was scaled to have a total variance 32 times larger than the RNA-seq data (Figure 2D).

### Coupled factors improve the accuracy of discerning MRSA bacteremia outcomes

We next sought to evaluate the extent to which CMTF-derived patterns could accurately distinguish MRSA PB vs. RB outcomes. We built a regularized logistic regression classifier to assign persistence outcome from the CMTF components and compared its performance to those built with a single data source (Figure 3A/B). As elevated IL-10 is associated with PB (4, 5, 16), we also constructed logistic regression models to assign persistence outcome from IL-10 measurements alone. We consistently observed that CMTF-derived factors could more accurately differentiate PB vs. RB outcomes. These results also revealed that the plasma cytokines were especially important to assignment accuracy, as CMTF performed better on patients having these measurements (Figure 3A). Note that, because each data source was not available for all patients, comparisons were made using the subset of patients having the respective data measurements. This is the reason for the performance of CMTF varying in each comparison. As further validation, we compared the prediction accuracy for a separate cohort that remained blinded during the model assembly process. While there was an expected decrease in accuracy overall, we again consistently observed that CMTF-derived factors were more effective at differentiating PB vs. RB outcomes (Figure 3C/D). Thus, data integration improved the accuracy of PB assignment, and CMTF-derived factors can have as high as 75% balanced accuracy when plasma cytokine samples are available (Figure 3A).

**Figure 3.**
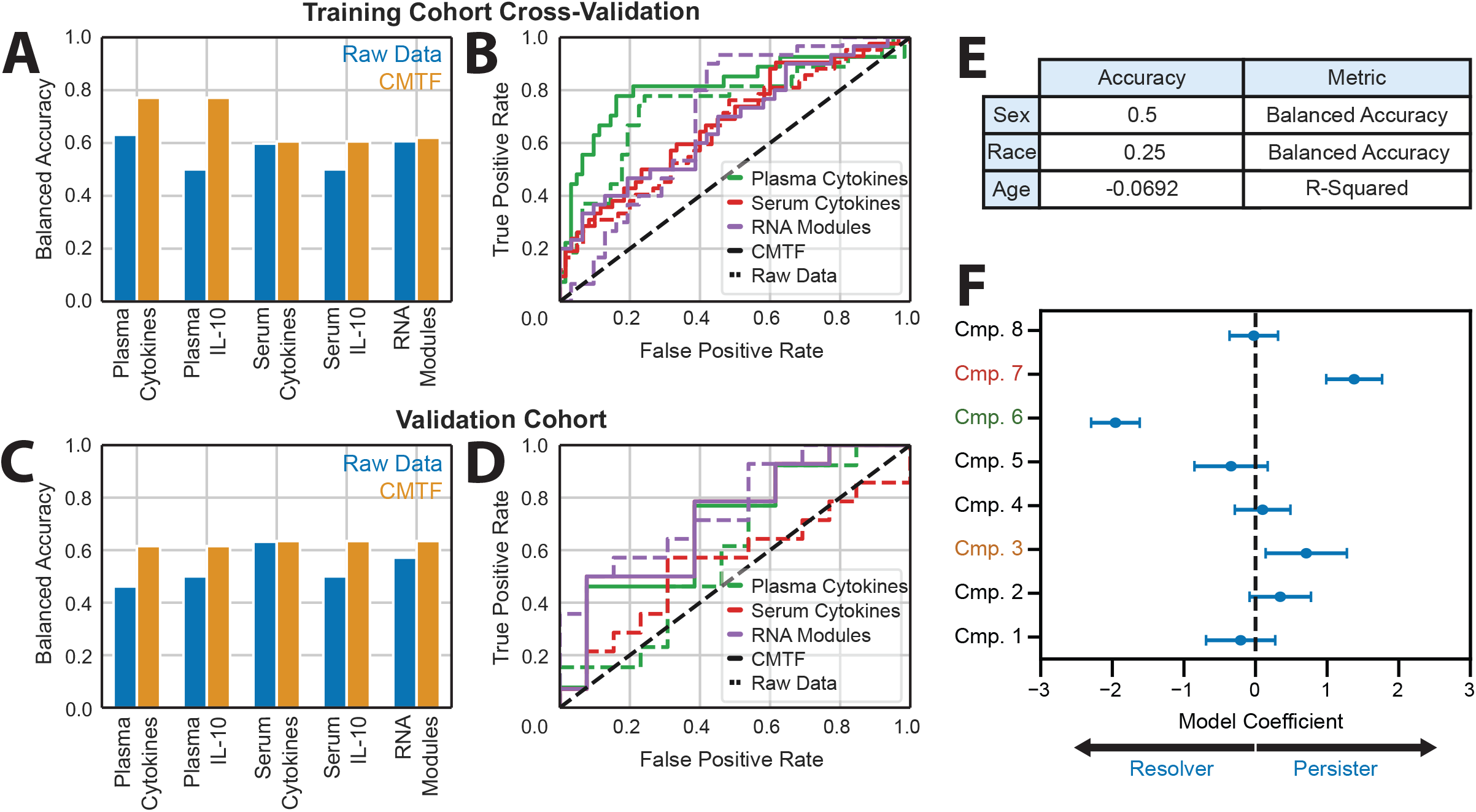
CMTF improves assignment accuracy of persistent MRSA bacteremia. A) Balanced accuracy in RB/PB assignment from models trained with CMTF components and raw data sources. Accuracy is evaluated using 10-fold cross-validation over the training cohort. B) Receiver operating characteristic curves for the models depicted in (A). C) Balanced accuracy in RB/PB assignment from models trained with CMTF or the raw data sources. Model accuracy is evaluated against a masked validation cohort following training against the training cohort. D) Receiver operating characteristic curves for models depicted in (C). E) CMTF model performance in assigning auxiliary demographics. F) Model coefficients assigned to each CMTF component. Dots and error bars depict the bootstrapping means and standard deviations of model coefficients, respectively.

We also sought to evaluate any relationships between the CMTF-derived components and patient demographic characteristics, specifically biological sex, age, and race (Figure 3E). MRSA susceptibility is known to vary with each of these parameters, and so we surmised that persistence may be influenced by factors that also vary with these characteristics (7). However, neither sex, age, nor race correlated better than chance, and there were no significant associations of the molecular components studied with these demographic features (Figure 3E).

A benefit of the logistic regression model is its ease of interpretation. As part of the fitting process, the logistic regression model assigns a coefficient to each CMTF component (Figure 3F). These coefficients indicate the relative impact of each CMTF component in MRSA PB vs. RB outcomes; more influential components have associated coefficients of greater magnitude. Additionally, the sign of the coefficient informs the directionality of the association; positive coefficients indicate an association with PB outcomes, while negative coefficients associate with RB outcomes. We quantified the uncertainty in each coefficient by bootstrapping the patient factors produced via CMTF (17). Notably, CMTF components 3 and 7 were strongly associated with the PB outcome, whereas component 6 was strongly associated with the RB outcome (Figure 3F). Overall, by magnitude, component 6 had the strongest correlation with MRSA bacteremia outcome.

### Coupled factors reveal conserved immunological responses in MRSA bacteremia

We plotted the composition of each CMTF component against the four factor dimensions (patient, cytokine, serum vs. plasma, and RNA expression module) to evaluate the relative biological significance of each component (Figure 4A–D). Factor components were scaled to have a dynamic range of −1 and 1. Next, gene enrichment analysis was performed to identify biological processes enriched within each gene module (Figure 4E).

**Figure 4.**
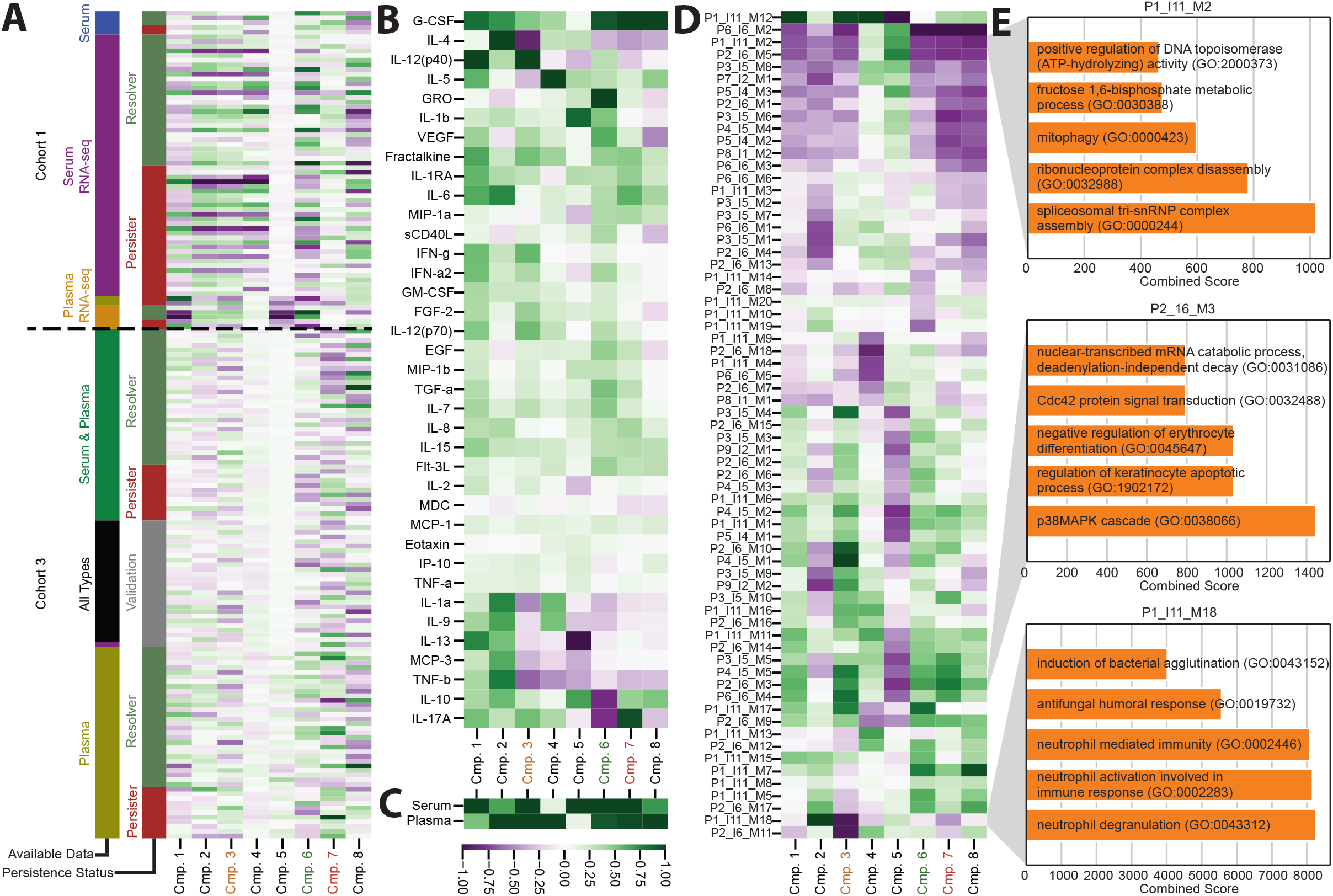
Components identify conserved patterns of MRSA immunologic response. A) Patient factors for each component. B) Component associations with each measured cytokine. C) Component associations with the two cytokine sources: plasma and serum. D) Component associations with each gene module. E) Selected enrichment analysis results for modules with high association to a persistence-associated component.

These factor matrices can be interpreted in two primary ways. To interpret the biological significance of a particular CMTF component, one can evaluate its composition across every factor matrix. For instance, component 3 corresponds to upregulation of IL-12(p40) and downregulation of IL-4 (Figure 4B) across both plasma and serum sources (Figure 4C). Component 3 also correlates with the downregulation of module P1_I11_M18 (Figure 4D) that enriches for gene sets involving neutrophil activation (Figure 4E). In parallel, one can also compare the differences between components and the variance they explain by examining each individual factor matrix. Components 6 and 7, for example, are mostly similar in their associations to RNA profiles (Figure 4D) and serum and plasma cytokine sources (Figure 4C) but are distinct in their association with specific cytokines (Figure 4B).

Figure 3F identifies components 3, 6, and 7 as important correlates of persistent MRSA bacteremia outcome. As shown in Figure 4, component 7, which correlates with PB outcomes, corresponds to upregulation of G-CSF, IL-17A, and the gene module P2_I6_M3. Enrichment analyses of P2_I6_M3 identifies significant enrichment of gene sets involving regulation of cellular apoptosis and differentiation. Component 6, which associates with RB outcomes, demonstrates some similarities to component 7 as it has similar RNA-seq associations and corresponds to G-CSF upregulation. Component 6, unlike component 7, also associates with the upregulation of GRO and the downregulation IL-10 and IL-17A.

The remaining CMTF components (1, 2, 4, 5, and 8) identify biological processes across RNA-seq and cytokine measurements that do not strongly associate with persistent MRSA bacteremia outcome. Of these components, we find that component 1 associates strongly with IL-12(p40) and gene module P1_I11_M12 upregulation; enrichment analysis of P1_I11_M12 was inconclusive, but we find genes relating to monocyte activation (see Supplementary Data). Component 2 corresponds with the upregulation of IL-4 and module P1_I11_M18 that enriches for gene sets involving neutrophil activation and immunity, suggesting that this component corresponds to a neutrophil activation pathway. Components 4 and 5, similarly to component 1, relate to P1_I11_M12 upregulation; these components are unique in their cytokine signatures, however, as they associate with strong IL-5 upregulation (component 4) and IL-13 downregulation (component 5). Finally, like components 6 and 7, component 8 associates strongly with G-CSF upregulation. Component 8, however, corresponds to both P6_I6_M2 downregulation and P1_I11_M7 upregulation that enrich for secretory granule lumens and cardiac tissue morphogenesis, respectively (refer to the enrichment results under *Data and Materials Availability*).

Components 1, 2, 4, and 5 appear to be primarily influenced by batch effects. Figure 4A shows that the magnitudes of these components vary greatly across cohorts, suggesting that these components identify signals with cohort-to-cohort differences. Dimensionality reduction techniques, including tensor factorization, are routinely used for correcting batch effects, and these batch-associated components help to capture and “remove” patterns associated to batch effects so that the remaining components may better associate with persistence-related mechanisms.

### A reduced model reveals heterogeneity in persistent MRSA bacteremia outcomes

Given that only a subset of components was strongly associated with MRSA persistence, we next sought to test whether a reduced model using few components could be equally associative with persistence as the full model. To do so, we fit support vector machine (SVM) classification models using pairings of components 3, 6, and 7 given their high associations with persistence in Figure 3F. We compared these reduced SVM models to a logistic regression model that uses all eight CMTF components. In contrast to our earlier analysis, we included both cohorts of patients in this analysis, examining all 177 patients. We observed that an SVM model trained with only components 6 (upregulation of G-CSF, GRO, and granulocyte activation and downregulation of IL-10, IL-17A) and 7 (upregulation of G-CSF, IL-17A, and granulocyte recruitment and downregulation of granulocyte activation) had equal balanced classification accuracy to the model built with all components (Figure 5A–B), supporting the notion that components 6 and 7 are informative of bacteremia persistence outcome on their own.

**Figure 5.**
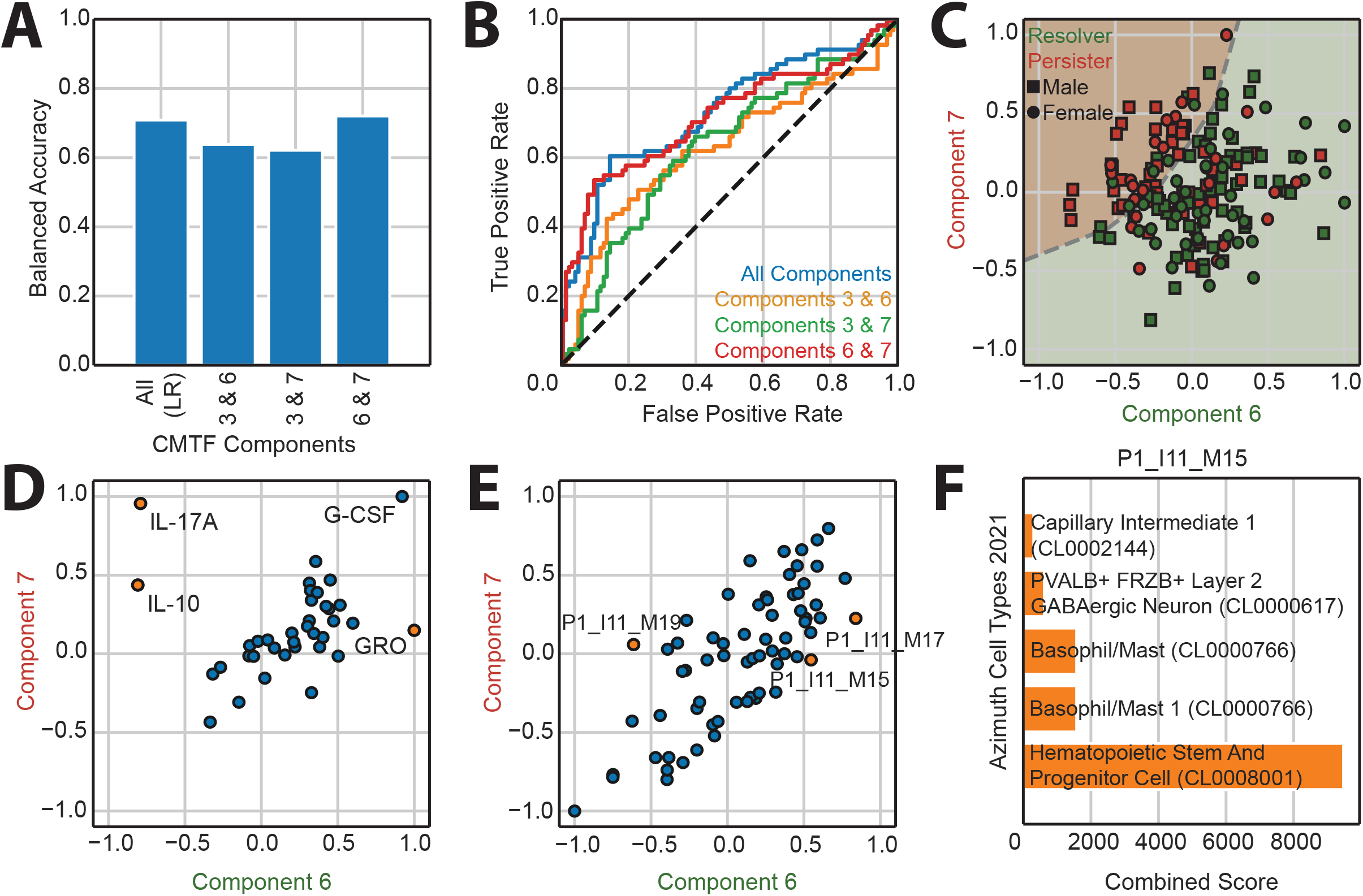
A reduced model visualizes heterogeneity in persistent MRSA bacteremia outcomes. A) Balanced accuracy scores of support vector machine classification models trained with subsets of CMTF components. Accuracy is evaluated using 10-fold cross-validation over all patients. B) Receiver operating characteristic curves for models depicted in (A). C) Patient factor values for CMTF components 6 and 7. The dashed line represents the persistence decision boundary of the SVM model. Purple and red shading indicate persister- and resolver-predicted patients, respectively. D) Cytokine factor values for CMTF components 6 and 7. Cytokine factors with large absolute differences between components 6 and 7 are highlighted in orange. E) RNA module factor values for CMTF components 6 and 7. RNA module factors with large absolute differences between components 6 and 7 are highlighted in orange. F) Enrichment analysis results for module P1_I11_M15.

With the reduced model indicating the independent value of components 6 and 7, we then plotted the patients within this reduced space (Figure 5C). Here, we find a clear subset of patients who experienced persistent MRSA bacteremia in the top-left that the model identifies as PB. This relationship indicates that negative component 6 values coupled with positive component 7 values are strongly associated with PB; that is, decreases in RB-associated component 6 in conjunction with increases in PB-associated component 7 associates with PB.

The combined importance of components 6 and 7 was surprising because they share many factor associations (Figure 4). To examine the distinction between components 6 and 7 more closely, we plotted the cytokine (Figure 5D) and RNA module (Figure 5E) factor values of components 6 and 7 against each other. We find that many cytokines demonstrate similar factor values across components 6 and 7. However, IL-17A and IL-10 are relatively much lower in component 6 while GRO is much higher. Likewise, the RNA module factors are also similar between components 6 and 7 except for P1_I11_M15 and P1_I11_M17, both of which are more positive in component 6, and P1_I11_M19, which is lower in component 6. Enrichment analyses of P1_I11_M15 shows significant enrichment for hematopoietic stem cell generation (Figure 5F). Enrichment analyses for P1_I11_M17 and P1_I11_M19 failed to find significantly enriched gene sets. However, these gene modules do contain fibroblast growth factor 23 (P1_I11_M17) and cytokine receptor expression (P1_I11_M19; see TableS1 under *Supplementary Data*).

## Discussion

Here, we demonstrate that CMTF can improve our understanding of host immune responses correlated with persistence outcomes in human MRSA bacteremia. This insight is gained by characterizing the host immune response through the integration of multiple data types. We find that CMTF captures over 70% of the variation observed across clinical RNA expression and proteomic measurements in just 8 components, that these components strongly associate with MRSA persistence (Figure 2), and that the components more accurately predict persistence outcome than RNA expression or proteomic measurements alone (Figure 3). Our prediction model indicates each component’s association to persistence, while CMTF supplies information about how each component relates to the individual measurements. Consequently, we can biologically interpret these components to better understand the underlying immunologic mechanisms (Figure 4). We find a subset of components are equally associated with persistence to our full model (Figure 5). Examination of this component subset more precisely delineates heterogeneity in immune responses to MRSA bacteremia. We have summarized this observed heterogeneity in immune responses and highlight the key immunological signatures associated with PB and RB outcomes in Figure 6.

**Figure 6.**
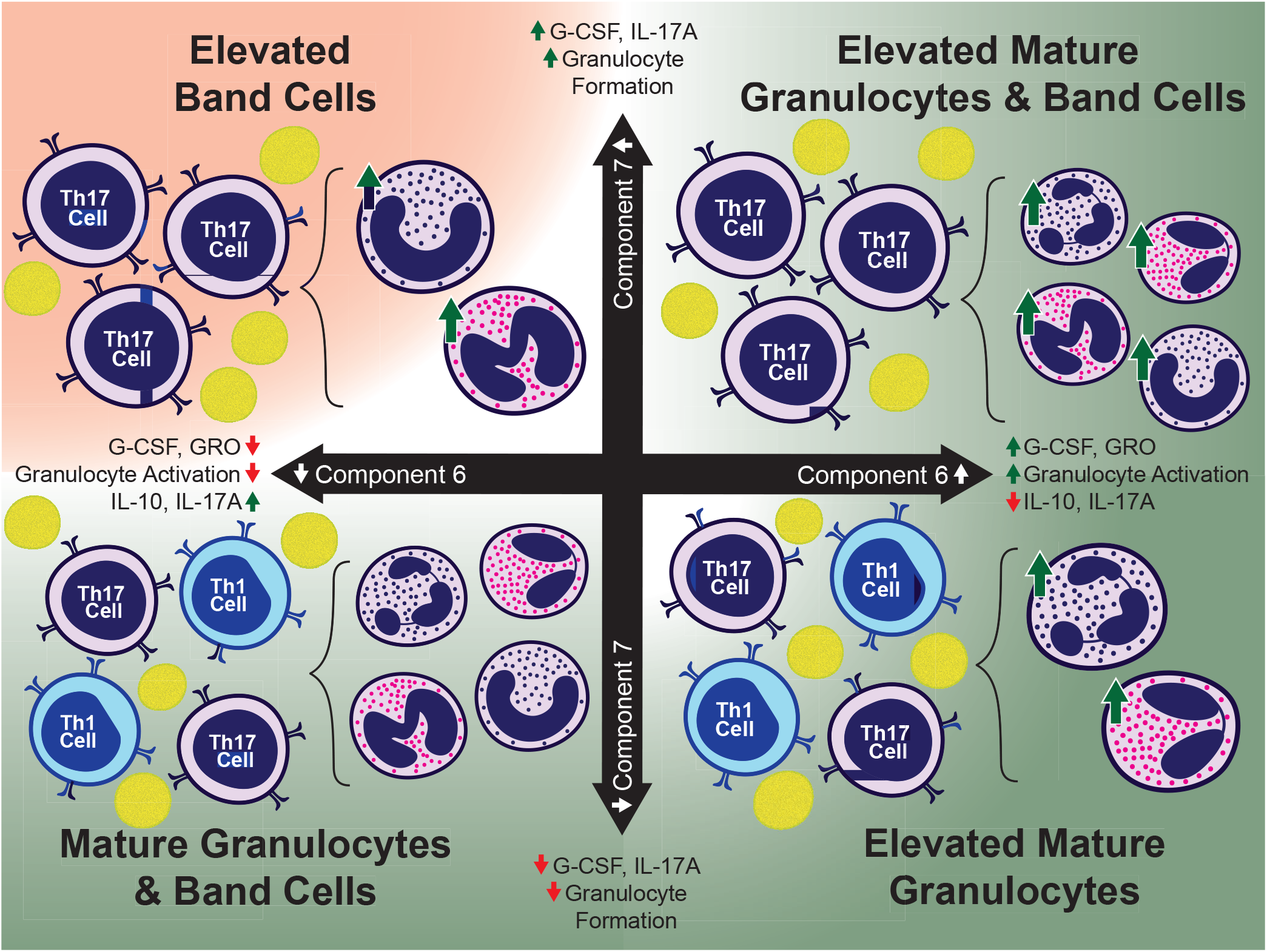
CMTF emphasizes role of T-helper cell polarization in granulocyte formation and in bacteremia persistence. Illustration depicting immunological signatures associated with combinations of component 6 and 7. Red shading indicates signatures associated with PB; green shading indicates association with RB.

In integrating the serum and plasma cytokine with whole blood RNA-seq measurements, we found that the plasma cytokine measurements were most important for evaluating persistence (Figure 3). While the integration of data types leads to optimal persistence assignment accuracy, the importance of the cytokine measurements over RNA-seq suggests that the former is more directly indicative of MRSA persistence. Additionally, while previous studies have found associations between sex, race, and age with persistent MRSA bacteremia (7), we found that the CMTF components are not associative with these auxiliary demographics. This perhaps suggests that our molecularly derived patterns are associated with persistence through independent means.

The observed differences in cytokine profiles typically generated by distinct CD4+ T helper cell polarization in response to infection implies T cell responses are integral to persisting versus resolving outcomes in MRSA bacteremia. This concept is consistent with findings regarding epigenetic correlates of such outcomes (4). In the present study, further interrogation of the cytokine and gene expression factors relevant to each component also support and extend existing literature in MRSA persistence mechanisms. Component 3 associates strongly with IL-12 (p40) upregulation that typically associates with antigen-presenting cell response early in the course of infection and subsequent downregulation of IL-4 and IL-10 (18–20). In contrast, component 3 also correlated with the downregulation of IL-4 and gene sets enriched in neutrophil activation. Neutrophils are essential for phagocytosis in the innate immune response via the Th17 pathway (21), and MRSA has well-documented mechanisms of avoiding neutrophil recognition (22). Additionally, IL-4 and IL-12 (p70) are both implicated in B cell maturation (23–25); as component 3 is associated with the downregulation of IL-4 and the upregulation of an IL-12 (p70) regulatory counterpart, IL-12 (p40), this component also potentially corresponds to a process that suppresses B cell maturation and signaling. This mechanism is also synergistic with neutrophil suppression as neutrophils, B cells, and IL-12 (p70) have documented cross-talk mechanisms in response to inflammation (25). Alternatively, IL-12 (p40) upregulation could correspond to an increase in IL-23, a pro-inflammatory, Th17-promoting cytokine that includes an IL-12 (p40) subunit (26). IL-23 promotion of inflammation and Th17 cells has been linked to pathogenic inflammation (26, 27), suggesting that, overall, component 3 may correspond to a non-protective inflammation mechanism in response to MRSA bacteremia.

Intriguingly, components 6 and 7 demonstrate some similar cytokine and RNA expression signatures but have opposite associations with persistence outcome. Both components strongly associate with G-CSF upregulation, which promotes neutrophil production and maturation (28), suggesting both components involve increases in neutrophil production. Component 7, however, also strongly associates with IL-17A upregulation whereas component 6 associates with IL-17A downregulation. Elevated levels of G-CSF and IL-17A are linked to neutrophilia (4, 29), suggesting that component 7 may correspond to a pathogenic neutrophil production mechanism. Component 6, conversely, corresponds to increased GRO which associates with improved neutrophil recruitment and infiltration (30), suggesting that component 6 associates with both improved neutrophil production and recruitment to sites of infection that may hematogenously see ongoing bacteremia. Component 6 also associates with the downregulation of IL-10, an immune-suppressing cytokine known to be aberrantly induced by MRSA (4, 5, 16), implicating the downregulation of IL-10 production in clearance of bacteremia in context of vancomycin therapy as in the current study cohort. In contrast, component 7 associates with the upregulation of gene sets associated with the negative regulation of erythrocyte differentiation, suggesting that component 7 corresponds to an immunosuppressive process that prevents immune cell differentiation. A recent study found that persistent MRSA outcomes correlate with suppressed neutrophil maturation, suggesting that component 7 may indeed identify a persister-associated mechanism that involves the proliferation of immature immune cells or their non-protective maturation (4). Altogether, these differences between components 6 and 7 highlight that immune cell maturation, differentiation, and trafficking are critical in resolving MRSA infections, and that immune cell production alone can be either persister- or resolver-associated. The key concept is that outcome of MRSA bacteremia is defined through a balance of these immune responses.

More broadly, these results find that a comprehensive view of cytokine and RNA expression data improves our understanding of the immune response to MRSA. Integrating data types to define molecular patterns of immunologic response provides dual benefits: Firstly, interpretation of the resulting patterns is made easier through a broader view of the involved molecular factors. Additionally, immune response patterns, especially with data reduction in tensor form, are more precisely defined through more effective dimensionality reduction. We find that the mechanisms of persistent MRSA bacteremia manifest over multiple biological modalities, and that tensor factorization methods can recognize patterns across modalities to identify these mechanisms of MRSA persistence. These results demonstrate the importance of multi-omics data profiling and integration in characterizing the human immune response, with coupled tensor factorization as a powerful tool for providing interpretable data integration.

## Materials and Methods

### Patients and sample collection

This case-controlled study consisted of 177 SAB patients (71 PB and 106 RB) propensity matched by sex, race, age, hemodialysis status, type I diabetes, and presence of an implantable device. Details of clinical characteristics of study cohort are presented in Table 1. Of the 177 patients, 129 had serum cytokine measurements, 115 had plasma cytokine measurements, and 88 had RNA-seq measurements. SAB cases were evaluated and consented for enrolment in the S. aureus Bacteremia Group (SABG) biorepository at Duke University Medical Centre (DUMC).

**Table 1.**
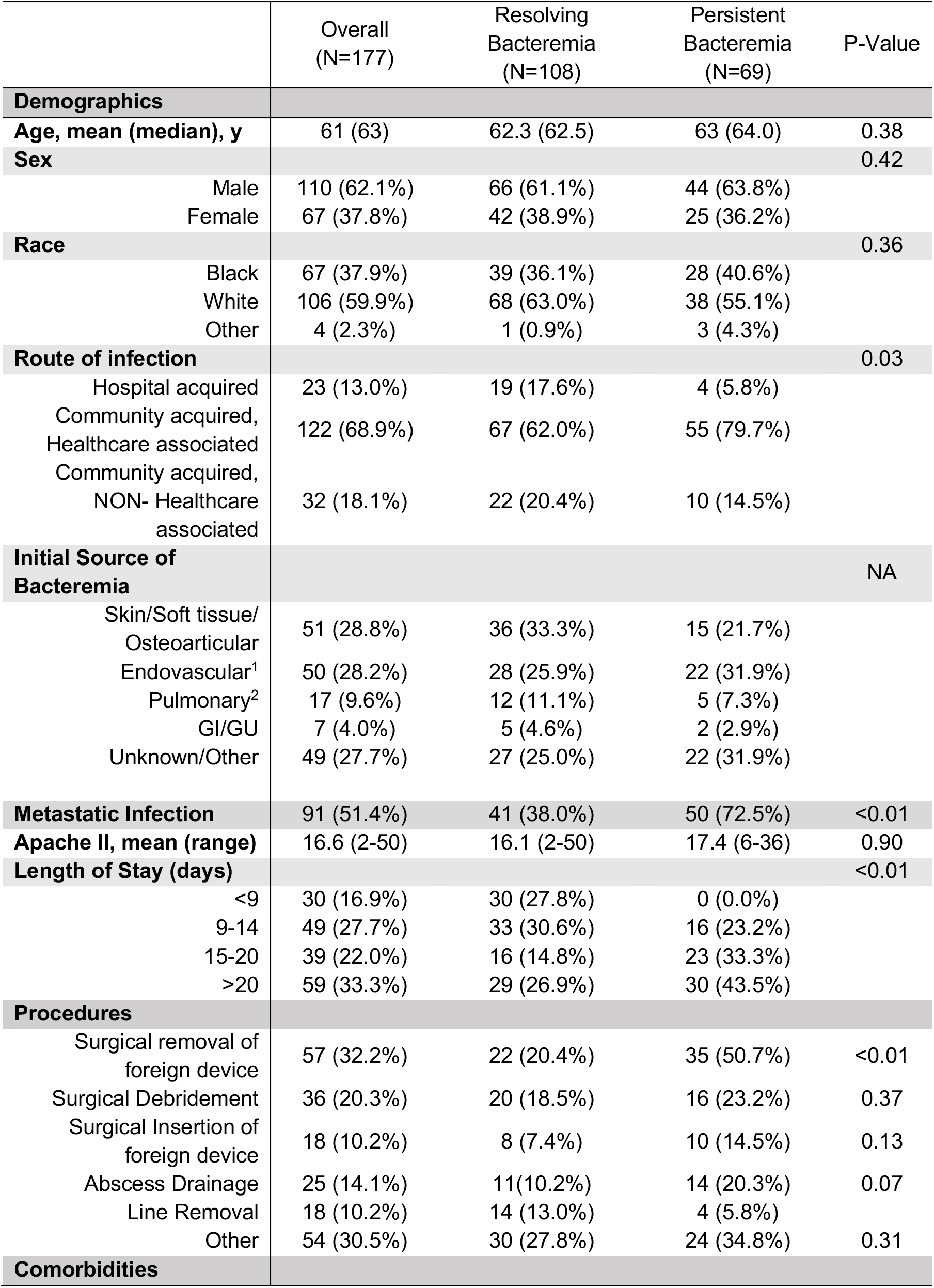
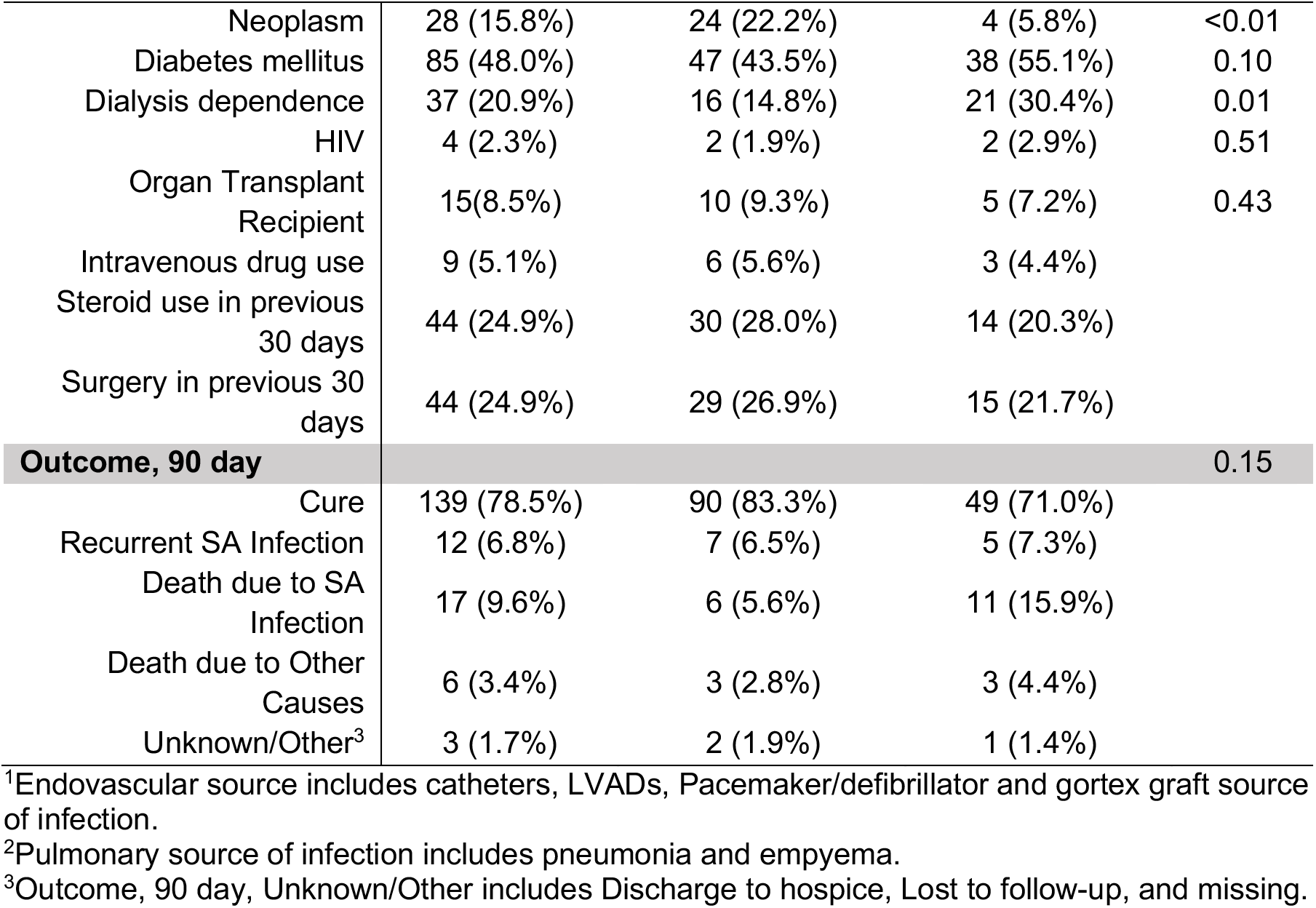
Demographic and Clinical Characteristics of the Patients with Resolving and Persistent Bacteremia.

Plasma and/or sera and whole blood Paxgene samples were collected at time of diagnosis of MRSA infection and stored in the SABG biorepository. Cases for the current study were carefully selected based on the following inclusion criteria: laboratory confirmed MRSA bacteremia; received appropriate vancomycin therapy; enrolled in the SABG study between 2007 and 2017 (to ensure contemporary medical practices) and had available serum or plasma samples. PB was defined as patients had continuous MRSA positive blood cultures for at least 5 days after vancomycin antibiotic treatment (5); while RB patients had initial blood cultures that were positive for MRSA, but the subsequent blood cultures were negative.

### Molecular analysis

#### Luminex-based cytokine measurement

Human 38-plex magnetic cytokine/chemokine kits (EMD Millipore, HCYTMAG-60K-PX38) were used per manufacturer’s instructions. The panel includes IL-1RA, IL-10, IL-1α, IL-1β, IL-6, IFN-α2, TNF-β, TNF-α, sCD40L, IL-12p40, IFN-γ, IL-12/IL-12p70, IL-4, IL-5, IL-13, IL-9, IL-17A, GRO/CXCL1, IL-8/CXCL8, eotaxin/CCL11, MDC/CCL22, fractalkine/CX3CL1, IP-10/CXCL10, MCP-1/CCL2, MCP-3/CCL7, MIP-1α/CCL3, MIP-1β/CCL4, IL-2, IL-7, IL-15, GM-CSF, Flt-3L/CD135, G-CSF, IL-3, EGF, FGF-2, TGF-β, and VEGF. Fluorescence was quantified using a Luminex 200TM instrument. Cytokine/chemokine concentrations were calculated using Milliplex Analyst software version 4.2 (EMD Millipore). Luminex assay and analysis were performed by the UCLA Immune Assessment Core.

#### RNA sequencing, mapping, quantification, and quality control

Total RNA was isolated with Qiagen RNA Blood kit, and quality control was performed with Nanodrop 8000 and Agilent Bioanalyzer 2100. Globin RNA was removed with Life Technologies GLOBINCLEAR (human) kit. Libraries for RNA-Seq were prepared with KAPA Stranded mRNA-Seq Kit. The workflow consists of mRNA enrichment, cDNA generation, and end repair to generate blunt ends, A-tailing, adaptor ligation and PCR amplification. Different adaptors were used for multiplexing samples in one lane. Sequencing was performed on Illumina Hiseq3000 for a single read 50 run. Each sample generated an average of 15 million reads. Data quality check was done on Illumina SAV. Demultiplexing was performed with Illumina Bcl2fastq2 v 2.17 program.

### Computational Analysis

#### Cytokine Normalization

Prior to analysis, cytokine data were separated into two matrices including the serum and plasma samples each. These cytokine measurements included available data from both cohorts. For both sets of cytokine measurements, values above or below the limit of detections were set to be equal to those limits. IL-12p70 had an unusually low limit within cohort 1, and so values were set to the lowest measured value to minimize biased results during normalization. As each cytokine may span a different range of values, measurements were first log transformed and then mean-centered for each cytokine across all patients. Finally, because tensor factorization attempts to explain variation among the data, we divided each matrix by its standard deviation to ensure equal overall variance.

#### RNA Processing

Gene expression counts were converted to transcripts per million (TPM). Measured genes with an average TPM below 1 were removed. Genes then were grouped into modules through Weighted Gene Correlation Network Analysis (WGCNA) (15); modules were determined using the TPM of each gene across all patients. Each gene appears in at most one module, and a cut-off of kME >0.8 was used to define module membership. The arithmetic mean TPM was calculated for every module for each patient, resulting in a matrix of mean module TPM for each patient. Prior to further analysis, the mean module expression is mean-centered and variance-scaled for each module across all patients.

#### Enrichment Analysis

Enrichment analyses were performed with Enrichr to interpret the biological significance of the gene modules. For each module, the genes contained within the module were compared to the Gene Ontology’s Biological Process 2021 library of gene sets. Both the (1) p-value via Fisher’s Exact Test and (2) combined score via Enrichr’s rank-correction test were derived for each gene set with overlap to a module. Only gene sets with a p-value < 0.05 were considered significantly enriched; for modules with multiple significantly enriched gene sets, gene sets with the largest combined scores were considered most significantly enriched.

#### Coupled Matrix-Tensor Factorization

We separated the data into a three-mode tensor organized by patient, cytokine, and measurement source (serum or plasma) coupled with a matrix of RNA modules for each patient (Figure 1B). Tensor operations were performed using Tensorly (31). Here, we reduced the molecular measurements into the sum of *R* Kruskal-formatted components:

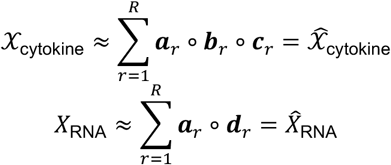

where ***a**_r_*, ***b**_r_*, ***c**_r_*, and ***d**_r_* are vectors indicating variation along the patient, cytokine species, cytokine sources (serum or plasma), and RNA module modes, respectively, and “o” indicates vector outer product. Concatenating all *R* vectors for each mode, we have their factor respective matrices, *A, B, C*, and *D*.

As we have reported elsewhere (13), tensor factorization was performed via an alternating censored least squares method (9). Each factor matrix was first initialized with imputed singular value decomposition of the unfolded tensor along its respective mode. With each iteration, leastsquares solving is performed separately for each mode with the missing values ignored. The cytokine factor matrix (*B*) is updated to the least-squares solution of the Khatri-Rao product (indicated by “ʘ”) of the cytokine source (*C*) and patient (*A*) factors and the cytokine tensor unfolded along the cytokine mode (*X*_cytokine, (2)_)

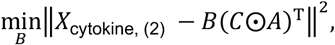

and the cytokines source matrix is updated in a similar fashion, with the unfolding performed along the source mode (*X*_cytokine, (3)_)

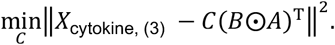

For the RNA modules matrix, the RNA factor matrix (D) is updated to the least-squares solution of the patient factors and RNA modules matrix

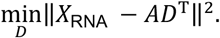

Finally, to enforce that the patient factors explain the variance across both datasets, the unfolded cytokine tensor is concatenated with the RNA module matrix

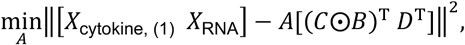

where *X*_cytokine, (1)_) indicates the tensor unfolding of 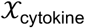 along its patient mode, and “[]” indicates the concatenation of two matrices within the bracket. Similarly, the Khatri-Rao product of the cytokine (*B*) and cytokine source (*C*) factors is concatenated with the RNA module factors (*D*) and the least-squares solution is derived using these concatenated matrices, leading to patient factors that minimize squared error overall. Iterations were repeated until the variance explained (R2X) improved by less than 1×10^-6^ between iterations.

#### Reconstruction Fidelity

To evaluate the fidelity of our factorization, we calculate the percent variance explained, R2X. First, the total variance is derived as the sum of the Frobenius norms squared of the cytokine tensor and RNA module matrix: 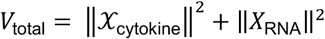. We then calculate the Frobenius norms squared of the difference between the cytokine tensor and RNA matrix and their reconstructed versions: 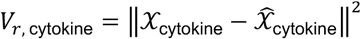 and 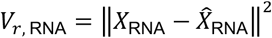. The variance explained is then calculated as

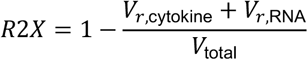

Missing values are ignored in all calculations.

#### Prediction Models

Following factorization, we use scikit-learn’s logistic regression classifier to predict persistence from CMTF’s patient factors (32). Data is regularized via elastic-net regularization with an 0.8 L1 ratio. Regularization strength is fitted via grid search using a stratified 10-fold cross-validation to evaluate prediction accuracy. This setup is also used for the single data source persistence prediction models (Figure 3A-D) and the CMTF race and sex prediction models (Figure 3E). Age is predicted using scikit-learn’s linear regression model (Figure 3E).

For the reduced models in Figure 5, we use scikit-learn’s support vector classification (SVC) model to predict persistence from pairs of components 3, 6, and 7. Data is regularized using L2 regularization with a regularization strength determined using the same scheme as the logistic regression model above.

### Data and materials availability

The code used to perform these analyses is available on GitHub at https://github.com/meyer-lab/tfac-mrsa. The cytokine and RNA-seq measurements are available in the GitHub repository at https://github.com/meyer-lab/tfac-mrsa/tree/main/tfac/data/mrsa. Enrichment results for each gene module are available at https://github.com/meyer-lab/tfac-mrsa/blob/main/tfac/data/mrsa/enrichment_results.zip. Gene memberships are available in **Dataset S1**.

## Acknowledgments

These studies were supported in part by NIH Grants U01-AI124319 (to M.R.Y.), R01-AI068804 (to V.G.F.), R33-AI111661 (to M.R.Y.), U01-AI124319 (to E.F.R.), and U19AI128913 (to E.F.R.).

## References

1. B. Borgundvaag, W. Ng, B. Rowe, K. Katz, EMERGency Department Emerging Infectious Disease Surveillance NeTwork (EMERGENT) Working Group, Prevalence of methicillin-resistant Staphylococcus aureus in skin and soft tissue infections in patients presenting to Canadian emergency departments. CJEM 15, 141–160 (2013).

2. M. L. Landrum, et al., Epidemiology of Staphylococcus aureus blood and skin and soft tissue infections in the US military health system, 2005-2010. JAMA 308, 50–59 (2012).

3. V. G. Fowler, Jr., et al., Persistent Bacteremia Due to Methicillin-Resistant *Staphylococcus aureus* Infection Is Associated with *agr* Dysfunction and Low-Level In Vitro Resistance to Thrombin-Induced Platelet Microbicidal Protein. J INFECT DIS 190, 1140–1149 (2004).

4. Chang Yu-Ling, et al., Human DNA methylation signatures differentiate persistent from resolving MRSA bacteremia. Proceedings of the National Academy of Sciences 118, e2000663118 (2021).

5. F. Mba Medie, et al., Genetic variation of DNA methyltransferase-3A contributes to protection against persistent MRSA bacteremia in patients. Proc Natl Acad Sci USA 116, 20087–20096 (2019).

6. L. C. Chan, et al., Protective immunity in recurrent *Staphylococcus aureus* infection reflects localized immune signatures and macrophage-conferred memory. Proc Natl Acad Sci USA 115, E11111–E11119 (2018).

7. S. I. Blot, K. H. Vandewoude, E. A. Hoste, F. A. Colardyn, Outcome and Attributable Mortality in Critically Ill Patients With Bacteremia Involving Methicillin-Susceptible and Methicillin-Resistant Staphylococcus aureus. Arch Intern Med 162, 2229 (2002).

8. V. Matzaraki, et al., Inflammatory Protein Profiles in Plasma of Candidaemia Patients and the Contribution of Host Genetics to Their Variability. Front. Immunol. 12, 662171 (2021).

9. T. G. Kolda, B. W. Bader, Tensor Decompositions and Applications. SIAM Review 51, 455–500 (2009).

10. T. Hastie, R. Tibshirani, J. H. Friedman, J. H. Friedman, The elements of statistical learning: data mining, inference, and prediction (Springer, 2009).

11. E. Acar, et al., Structure-revealing data fusion. BMC Bioinformatics 15, 239 (2014).

12. E. Acar, T. G. Kolda, D. M. Dunlavy, All-at-once Optimization for Coupled Matrix and Tensor Factorizations. arXiv:1105.3422 [physics, stat] (2011) (May 13, 2021).

13. Z. C. Tan, M. C. Murphy, H. S. Alpay, S. D. Taylor, A. S. Meyer, Tensor-structured decomposition improves systems serology analysis. Mol Syst Biol 17 (2021).

14. E. Acar, R. Bro, A. K. Smilde, Data fusion in metabolomics using coupled matrix and tensor factorizations. Proceedings of the IEEE 103, 1602–1620 (2015).

15. P. Langfelder, S. Horvath, WGCNA: an R package for weighted correlation network analysis. BMC Bioinformatics 9, 559 (2008).

16. J. Wang, G. Roderiquez, M. A. Norcross, Control of Adaptive Immune Responses by Staphylococcus aureus through IL-10, PD-L1 and TLR2. Scientific Reports 2, 606 (2012).

17. B. Efron, Nonparametric estimates of standard error: The jackknife, the bootstrap and other methods. Biometrika 68, 589–599 (1981).

18. D. Ashour, et al., IL-12 from endogenous cDC1, and not vaccine DC, is required for Th1 induction. JCI Insight 5 (2020).

19. n F. Koch, et al., High level IL-12 production by murine dendritic cells: upregulation via MHC class II and CD40 molecules and downregulation by IL-4 and IL-10. The Journal of experimental medicine 184, 741–746 (1996).

20. G. Liu, et al., Dendritic cell SIRT1–HIF1α axis programs the differentiation of CD4+ T cells through IL-12 and TGF-β1. Proceedings of the National Academy of Sciences 112, E957–E965 (2015).

21. M. R. Yeaman, et al., Mechanisms of NDV-3 vaccine efficacy in MRSA skin versus invasive infection. Proceedings of the National Academy of Sciences 111, E5555–E5563 (2014).

22. K. M. Rigby, F. R. DeLeo, Neutrophils in innate host defense against Staphylococcus aureus infections. Semin Immunopathol 34, 237–259 (2012).

23. Hsieh C S, Heimberger A B, Gold J S, O’Garra A, Murphy K M, Differential regulation of T helper phenotype development by interleukins 4 and 10 in an alpha beta T-cell-receptor transgenic system. Proceedings of the National Academy of Sciences 89, 6065–6069 (1992).

24. F. Ethuin, et al., Regulation of Interleukin 12 p40 and p70 Production by Blood and Alveolar Phagocytes During Severe Sepsis. Laboratory Investigation 83, 1353–1360 (2003).

25. A. Mantovani, M. A. Cassatella, C. Costantini, S. Jaillon, Neutrophils in the activation and regulation of innate and adaptive immunity. Nature Reviews Immunology 11, 519–531 (2011).

26. C. Tang, S. Chen, H. Qian, W. Huang, Interleukin-23: as a drug target for autoimmune inflammatory diseases. Immunology 135, 112–124 (2012).

27. G. R. Yannam, T. Gutti, L. Y. Poluektova, IL-23 in infections, inflammation, autoimmunity and cancer: possible role in HIV-1 and AIDS. J Neuroimmune Pharmacol 7, 95–112 (2012).

28. A. W. Roberts, G-CSF: a key regulator of neutrophil production, but that’s not all! Growth factors 23, 33–41 (2005).

29. S. B. Forlow, et al., Increased granulopoiesis through interleukin-17 and granulocyte colony-stimulating factor in leukocyte adhesion molecule-deficient mice. Blood 98, 3309–3314 (2001).

30. K. De Filippo, et al., Mast cell and macrophage chemokines CXCL1/CXCL2 control the early stage of neutrophil recruitment during tissue inflammation. Blood 121, 4930–4937 (2013).

31. J. Kossaifi, Y. Panagakis, A. Anandkumar, M. Pantic, TensorLy: Tensor Learning in Python. Journal of Machine Learning Research 20, 1–6 (2019).

32. F. Pedregosa, et al., Scikit-learn: Machine learning in Python. the Journal of machine Learning research 12, 2825–2830 (2011).

33. Z. Chitforoushzadeh, et al., TNF-insulin crosstalk at the transcription factor GATA6 is revealed by a model that links signaling and transcriptomic data tensors. Sci. Signal. 9, ra59–ra59 (2016).

34. X. Zhang, L. Li, Tensor Envelope Partial Least-Squares Regression. Technometrics 59, 426–436 (2017).

