## Supplemental Figures for "Cytokine-expression patterns reveal coordinated immunological programs associated with persistent MRSA bacteremia"

**This PDF file includes:**

Figures S1 to S2
Dataset S1

**Figure S1. Cytokine measurements vary by source and persistence status.** A) Pearson's correlation coefficient between plasma and serum cytokine measurements for each measured cytokine. B–C) Boxplots depicting normalized serum (B) and plasma (C) cytokine measurements. Boxplots for each cytokine are separated by persistence outcome.

**Figure S2. Correlations between cytokines.** Pearson's correlation coefficients between cytokine species for serum (A) and plasma (B) cytokine measurements.

**Dataset S1 (separate file). WGCNA Gene module memberships.** Columns include the Ensembl and Entrez gene names (if available) for each gene measured via RNA-seq. The kME metric defines the strength of a gene's membership to its module with higher scores indicating a stronger membership.

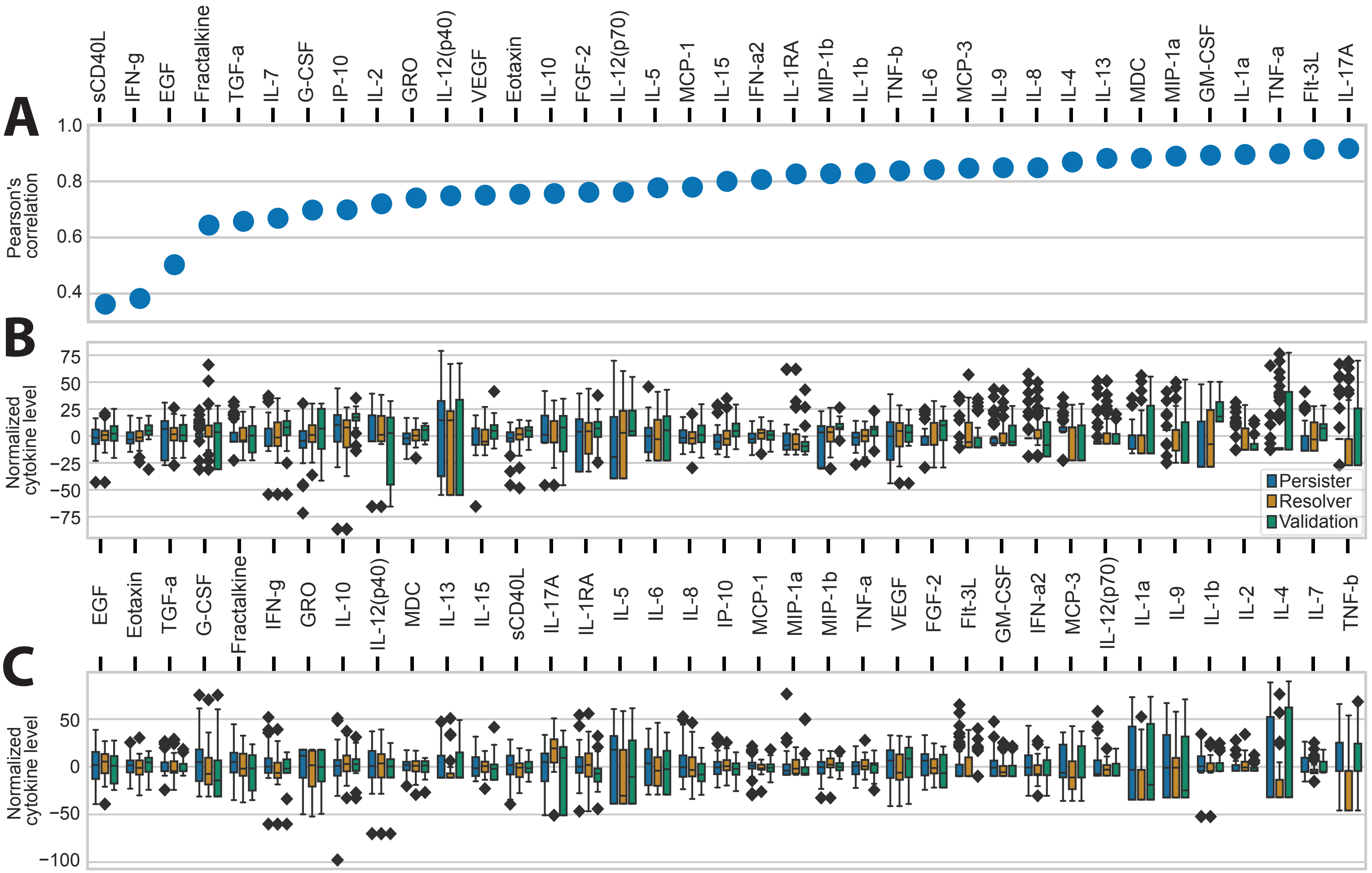
A
B
C
Pearson's correlation
Normalized cytokine level
Persister
Resolver
Validation
sCD40L
IFN-g
EGF
Fractalkine
TGF-a
IL-7
G-CSF
IP-10
IL-2
GRO
IL-12(p40)
VEGF
Eotaxin
IL-10
FGF-2
IL-12(p70)
IL-5
MCP-1
IL-15
IFN-a2
IL-1RA
MIP-1b
IL-1b
TNF-b
IL-6
MCP-3
IL-9
IL-8
IL-4
IL-13
MDC
MIP-1a
GM-CSF
IL-1a
TNF-a
Flt-3L
IL-17A

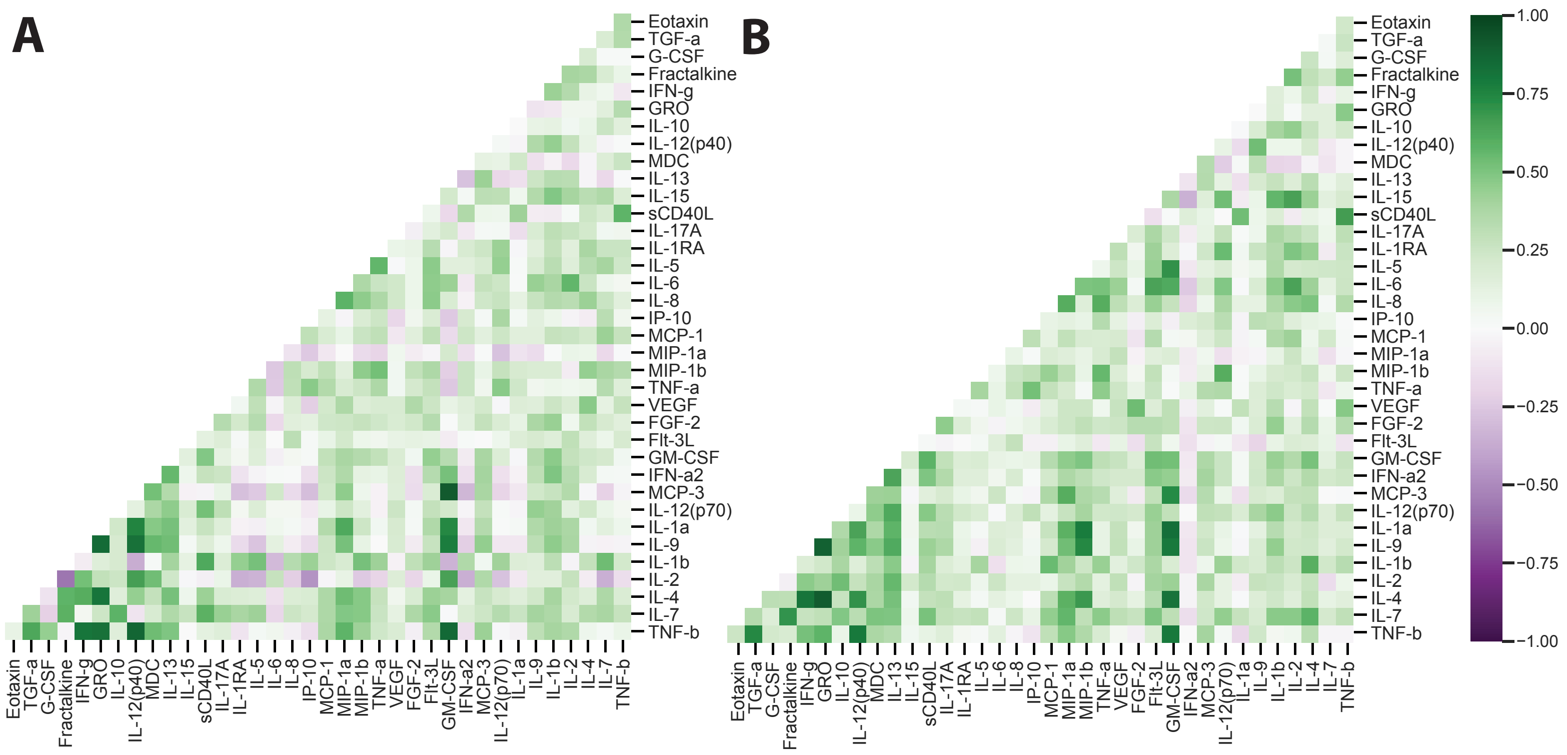

A
B
Eotaxin
TGF-a
G-CSF
Fractalkine
IFN-g
GRO
IL-10
IL-12(p40)
MDC
IL-13
IL-15
sCD40L
IL-17A
IL-1RA
IL-5
IL-6
IL-8
IP-10
MCP-1
MIP-1a
MIP-1b
TNF-a
VEGF
FGF-2
Flt-3L
GM-CSF
IFN-a2
MCP-3
IL-12(p70)
IL-1a
IL-9
IL-1b
IL-2
IL-4
IL-7
TNF-b
1.00
0.75
0.50
0.25
0.00
−0.25
−0.50
−0.75
−1.00
